# Inhibiting POLQ-mediated alternative NHEJ enhances CRISPR/Cas9 mediated precise genome editing in CHO cells

**DOI:** 10.1101/2022.12.09.519421

**Authors:** Chuanjie Wang, Ming Wang, Mengmeng Zhang, Yao Wang, Xinying Li, Chenghua Liu, Rongrong Fan, Yuanqiang Zheng, Beifen Shen, Zhaolin Sun, Jing Wang, Jiannan Feng

## Abstract

CRISPR/Cas9 mediated precise gene editing requires homology-directed repair (HDR), which occurs less frequently than non-homologous end-joining (NHEJ) including the canonical NHEJ and alternative NHEJ (Alt-EJ) in mammalian cells, especially in CHO cells that inherent resist HDR. To solve the above hurdle, here we for the first time show that knockout the DNA polymerase θ (POLθ), which is essential for Alt-EJ, significantly increases the knock-in efficiency by nearly forty-fold in CHO cells via eGFP reporter system and does not affect the normal growth and proliferation of cells. Meanwhile, even when transfecting simple circular, without negative element homologous template DNA donor and CRISPR/Cas9 plasmid to two different genomic sites, the knock-in rate of 4kb donor integration can still reach a mean of over 80% (29/36) and 2.7% (1/36) of the selected cell colonies in *POLQ*^-/-^ CHO cells, however, no positive knock-in cell colonies was obtained in wild-type CHO cells which respectively selected 62 cell colonies and 36 cell colonies. Furthermore, we show that *POLQ* promotes random integration in CHO cells. Finally, RNA-sequence analysis reveals not significant altered DNA repair, metabolism, apoptosis, and cell cycle in *POLQ*^-/-^ cells. These findings open a new target gene *POLQ* to overcome bottlenecks of the precision genome editing.

## INTRODUCTION

The advent of zinc-finger nuclease (ZFN), transcription activator-like effector nuclease (TALEN) and most recently CRISPR (clustered regularly interspaced short palindromic repeats)/Cas9 (CRISPR-associated protein 9) technologies has changed the landscape of gene targeting (1-6). With CRISPR-Cas9, a guide RNA with a spacer sequence complementary to the target DNA directs DNA cleavage by the Cas9 endonuclease (7-10). These customizable nucleases efficiently create double-strand breaks (DSBs) at the target locus in the genome.

Mammalian cells repair DSB by multiple pathways, including the error-prone canonical NHEJ (C-NHEJ) (11,12), alternative NHEJ (Alt-EJ) (13-15) and high-fidelity homologous recombination (HR) pathways (16-18). The C-NHEJ and Alt-EJ pathways proceed by ligation of DNA ends after they have been processed and result in imprecise indels (generally small insertions or deletions) to achieve efficient gene disruption (9,10,19-21). In contrast to end-joining pathways, homology dependent repair (HDR) using an exogenous DNA repair template supports precise genome editing which enables the introduction of minor sequence modifications or larger stretches of novel DNA (22-25). Importantly, precise genome editing is highly sought after in the of model systems engineering, cell production of recombinant therapeutic proteins, and gene therapy (6,26,27). However, in most mammalian cell types especially in CHO cells, HDR is significantly limited by the competing C-NHEJ and Alt-EJ mechanisms which are far more efficient and error prone (15,28,29). The low efficacy of HDR remains the bottleneck in the various application of the precise genome editing and has been a topic of intensive interest.

Recently, several groups have reported to increase HDR by four main technical strategies. The first is to inhibit C-NHEJ pathway by knockout the key NHEJ factors such as the DNA ligase IV or transient inhibition of key NHEJ factors such as Ku70, DNA ligase IV or the DNA-dependent protein kinase catalytic subunit, via short hairpin RNA knockdown, small-molecule inhibition such as Scr7 and NU7411 or proteolytic degradation, increased HDR in mammalian cell lines and animals (30-34). The second is to augment HDR pathway by small-molecule stimulation such as RS-1 (35), or overexpressing DNA repair proteins such as RAD51 (36) and phage single-stranded DNA-annealing proteins (SSAP) (37), or co-express RAD52 and dn53BP1 (38), or recruiting HDR-related repair proteins to Cas9 such as dominant-negative p53-binding protein 1 (dn53BP1), CtIP, mSA, RAD51, and Brex27 (39-44). The third is to control the cell cycle by cell synchronization (45,46), or restricting Cas9 expression to the S/G2/M portion of the cell cycle by fusing Cas9 to a domain of Geminin (47). The finally is to modification and optimization of repair template (48-51). However, most previously studies have found that the efficiency of HR improvement is not very high, mostly about 2-fold (35-44), although Scr7 improves HR efficiency up to 19-fold in MelJuSo cells using short fragment donor (33). However, this effect is obviously acting in a cell type dependent manner which it does not work on many mammalian cells (35,46,47,52-54), especially CHO cells that naturally resist HR (55). At the same time, it is obvious that the Alt-EJ pathway mediated by *POLQ*, another important competition pathway for HR, has been poorly studied (56,57). Although the latest report that dual inhibition of *POLQ* and *LIG4* can completely inhibit RI (58,59), but knock out *POLQ* alone did not affect HR. So, improving HR-mediated genome editing in mammalian cells thus remains a major challenge, especially in CHO cells, which are widely used in biopharmaceuticals and inherent resist HR (55).

Here, we for the first time reported that only knockout the A-family DNA polymerase θ (POLθ) increases the knock-in efficiency by nearly forty-fold in CHO cells, via enhanced green fluorescent protein (eGFP) reporter system and does not affect the normal growth and proliferation of cells. Meanwhile, when transfecting circular homologous template DNA without negative element and CRISPR/Cas9 plasmid at two different genomic sites such as *ROSA26* and *COSMC*, the knock-in rate of large DNA fragment integration can reach a mean of over 80% and 2.7% of the selected cell colonies in *POLQ*^-/-^ CHO cells, while the mean knock-in rate is 0% in wild-type CHO cells. Finally, we show that *POLQ* promotes random integration via cell selected assay and knockout *POLQ* not significant altered DNA repair, metabolism, apoptosis, and cell cycle by RNA-sequence.

## RESULTS

### Generation of *POLQ*^-/-^ CHO Cells by CRISPR/Cas9

We hypothesized that inhibiting Alt-EJ will increase the chance of knock-in events in the CRISPR/Cas9-mediated gene editing system. Therefore, we selected the *POLQ*, which were the important genes involved in Alt-EJ pathways, for targeting with the CRISPR/Cas9. To construct the CRISPR/Cas9 expression vector, the pX330 vector was used the backbone vector (67). To design the sgRNA of the *POLQ*, the previously reported CRISPy tool was used (68). As shown in **Supplementary Figure S1A**, three sgRNAs were designed against the *POLQ* exon 1. First, the activities of three sgRNAs in CHO cells were screened with T7EI assay. The three px330-sgRNAs vectors were transfected into the CHO cells, respectively. After 72 h, the genome of the sgRNA-treated CHO cells was extracted and analyzed. Using ImageJ software analysis, the percentage of indels generated at the *POLQ* loci was estimated to be 0 %, 38.4%, 33.9%, respectively (**Supplementary Figure S1B**). Subsequently, TA-cloning and a DNA sequencing analysis of the PCR amplicons corroborated the induction of CRISPR/Cas9-mediated mutations at the targeted site (**Supplementary Figure S1C**). px330-gRNA2 cleaved the target site with the greatest efficiency. Therefore, we used px330-gRNA2 in subsequent experiments. To knockout *POLQ* gene, px330-sgRNA2 vector was transfected into CHO cells. Two days after transfection, the single cell was sorted by FACS. Individual cell clones were isolated 5–7 days after FACS sorted, and the single clones were then expanded in culture, sequence analyzed and cryopreserved after a total of 12–14 days in culture. The indels generated at *POLQ* sgRNA target sites in single clones were analyzed by PCR and TA-clone. Finally, three *POLQ* ^-/-^ cell colonies were obtained from the 8 identified cell colonies (**Table 1**), the biallelic frame-shift mutations sequence was as shown in **Figure 1A**. The biallelic frame-shift mutations leaded to premature termination of translation, resulting in possible production of a peptide of different AA (**Figure 1B**). Furthermore, Western-blotting confirmed the successfully knockout of *POLQ* of the three CHO cell clone (**Figure 1C**). Meanwhile, the *POLQ* ^-/-^ cells grew normal compare with the WT cells (**Figure 1D**).

**Table 1.** Results of POLQ-knockout cell screening.

| Cell line | Isolated colonies | Positive cell colonies | Deletion Efficiency (%) | Double Deletion Efficiency (%) |
| --- | --- | --- | --- | --- |
| CHO-K1 | 280 | 47 | 16.7 (47/280) | 36.2 (17/47) |

**Figure 1.**
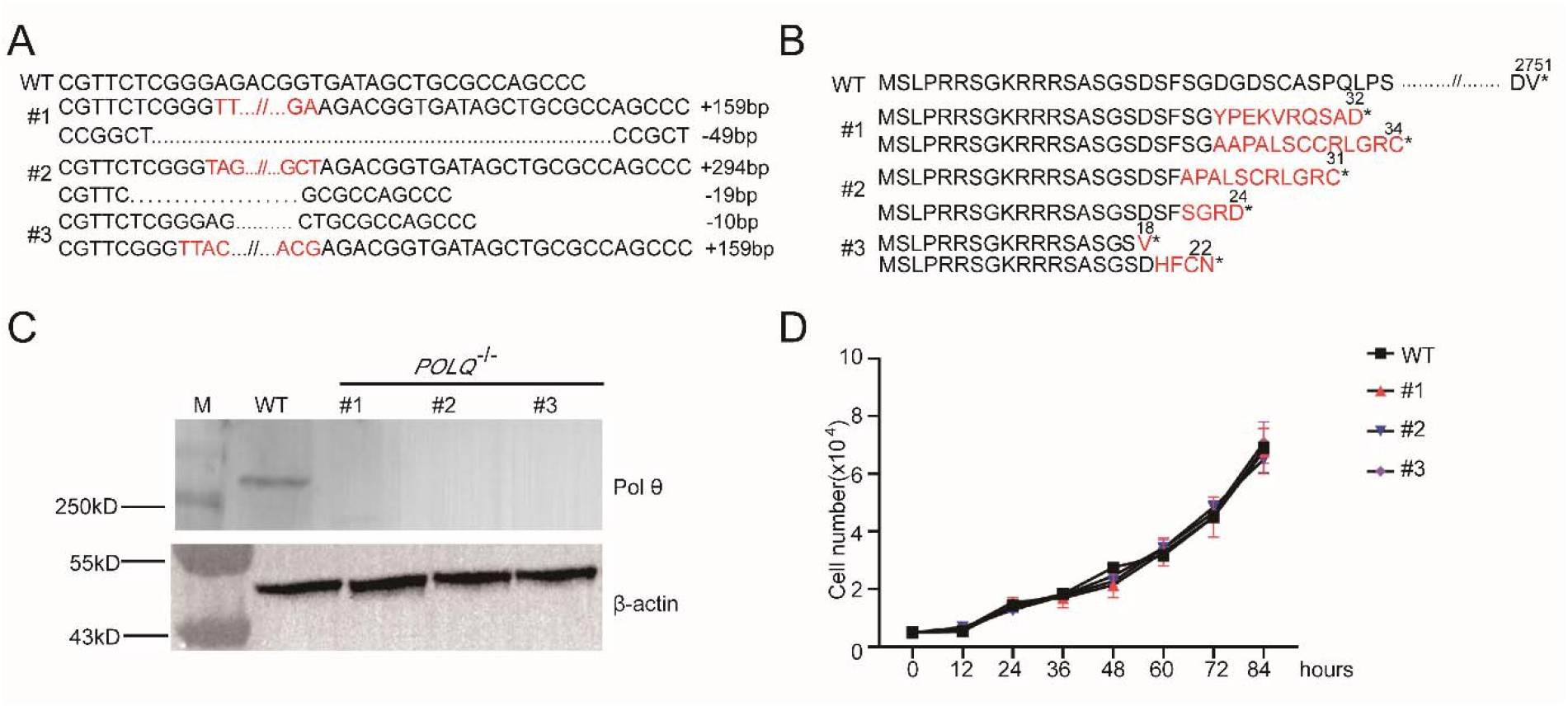
Generation and characterization of the *POLQ*^*−/−*^ knock-out CHO cells. **(A)** Sequences of the CRISPR-Cas9-induced mutations in *POLQ*^−/−^ cells. **(B)** The predicted changes to the protein resulting from *POLQ*^−/−^ cells. * represents the termination of protein translation. **(C)** Western blotting analysis to confirm the absence of Pol θ expression. WT, wild type cells; #1-#3, *POLQ*^−/−^ cell lines. β-actin was used as a reference. **(D)** Growth curves of the *POLQ*^−/−^ knock-out cell lines. Data shown are the mean ± SEM (n = 3).

### Loss of Pol θ robust increases HDR efficiency

To examine whether the loss of Pol θ can improve the HDR efficiency in the CHO cells, the green fluorescence protein reporter system was employed based on previously research(69,70). We aimed to fuse a p2A-EGFP reporter gene to intron of the *Rosa26* in *POLQ*^-/-^ and WT CHO cells. The resulting knock-in efficiencies are presented as percentages of GFP^+^ cells (**Figure 2A and 2B**). The sgRNA which target the intron of the *Rosa26* gene was used the recently reported study (**Table S1**). At 3 days after co-transfecting *POLQ*^-/-^ CHO cells and WT CHO cells with donor and the pX330-sgRNA plasmid, respectively. Interestingly, we observed a nearly 30-fold increase in the portion of GFP^+^ cells in *POLQ* ^-/-^ CHO cells compared to WT CHO cells (**Figure 2C and 2D**).

**Figure 2.**
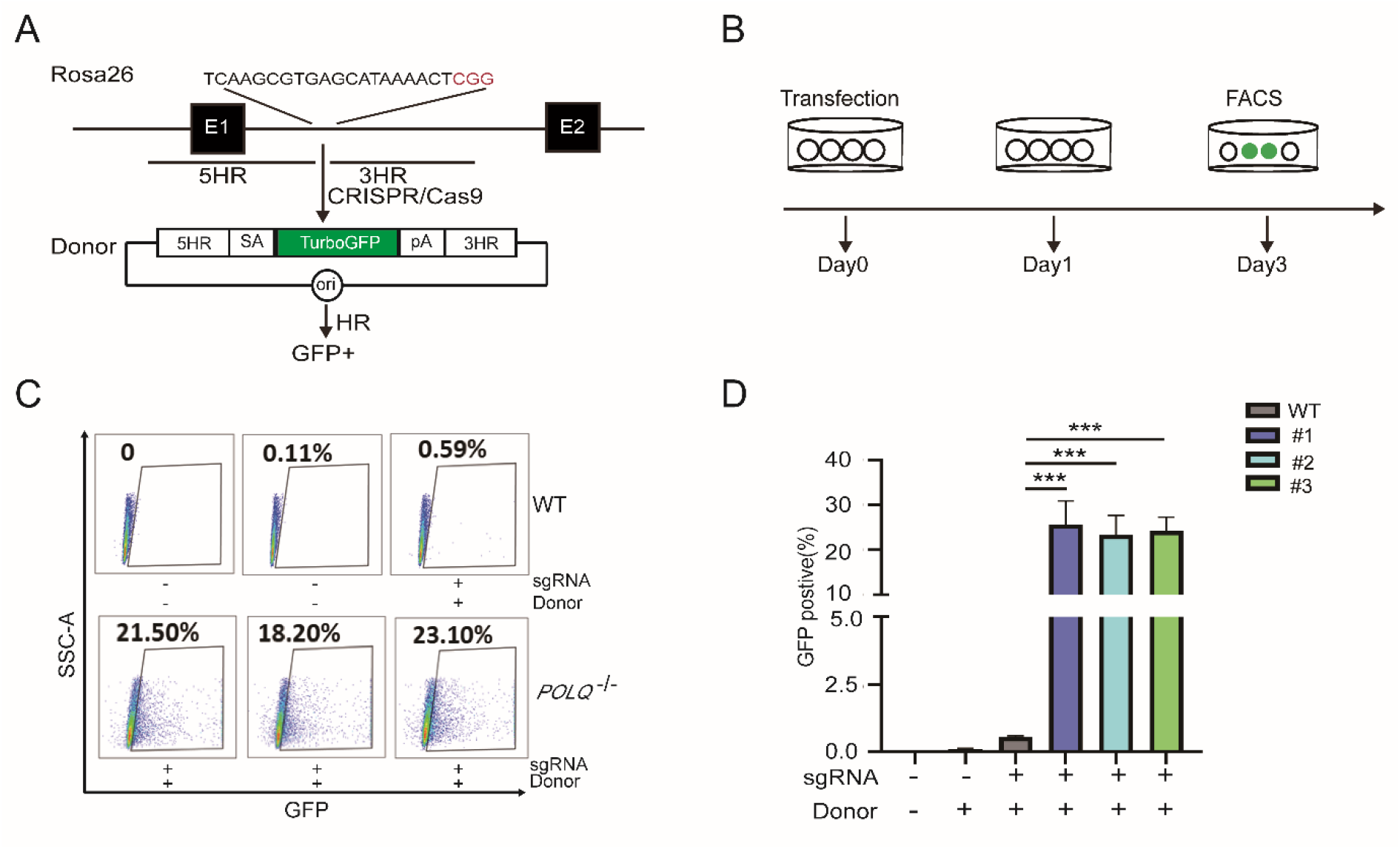
CRISPR-Cas9-stimulated GFP-gene targeting in *POLQ*^−/−^ CHO cells. **(A)** Strategy for insertion of a eGFP reporter gene into the *Rosa26* locus using CRISPR-Cas9 in CHO cells. The CRISPR/Cas9 target sequence (20-bp target and 3-bp PAM sequence (colored in red)) against the *Rosa26* locus is shown; in the Rosa26-EGFP targeting vector the GFP gene is flanked by *Rosa26* homology regions. **(B)** Schematic of gene-targeting assay in CHO cells to calculate GFP-targeting efficiency. Cells were cotransfected with linearized Rosa26-EGFP targeting vector and Rosa26-sgRNA, the percentage of GFP^+^ cells was analyzed by FACS 3 days after transfection. **(C)** FACS analysis for GFP^+^ cells in WT, *POLQ*^−/−^ CHO-KI cells. **(D)** Integration frequency of GFP in WT, *POLQ*^−/−^ CHO-KI cells, data shown are the mean ± SEM (n = 3).

### Highly efficient HDR at different locus using simple circular and without negative element donor and CRISPR/Cas9 in *POLQ*^-/-^ CHO cells

Meanwhile, to tested whether the high HR efficiency we observed in *POLQ*^-/-^ cells will be maintained in a more typical application scenario. We use a stable and integrated strategy for HR efficiency testing in two different locus and the simple circular and without negative element donor was used based on the above high HR efficiency (**Figure 3A** and **Supplementary Figure S2A**). For *Rosa26* locus, the simple donor pRosa26-HR target intron 1 of *Rosa26*, which included the 5’ homology arm, p2A-EGFP, the PGK-puro and 3’ homology arm. After co-transfected simple donor construct and pX330-sgRNA into *POLQ*^-/-^ and WT CHO cells (**Figure 3A**). After viable cells were selected using puro for approximately 14 days, surviving cell colonies were collected and then analyzed individually by PCR (**Figure 3B, 3C**). In total 36 cell colonies were identified and the 80% (25/36) efficiency of gene knockin was observed at the RosaA26 locus in *POLQ*^-/-^ CHO cells, while no positive knock-in cell colonies were obtained in wild-type CHO cells (0/65) (**Figure 3D**). DNA sequencing further confirmed the correctly targeted integration (**Figure 3E, 3F**). The *COSMC* locus, which was lower HR efficiency by previously reported, was also used to detect. In total 36 cell colonies were identified and the 2.8% (1/36) efficiency of gene knockin was observed at the *COSMC* locus in *POLQ*^-/-^ CHO cells, however, no positive knock-in cell colonies were obtained in WT CHO cells (0/36) (**Supplementary Figure S2B-E**). Taken together, the above results conducted in CHO cells suggest that loss of Pol θ leads to significant HDR.

**Figure 3.**
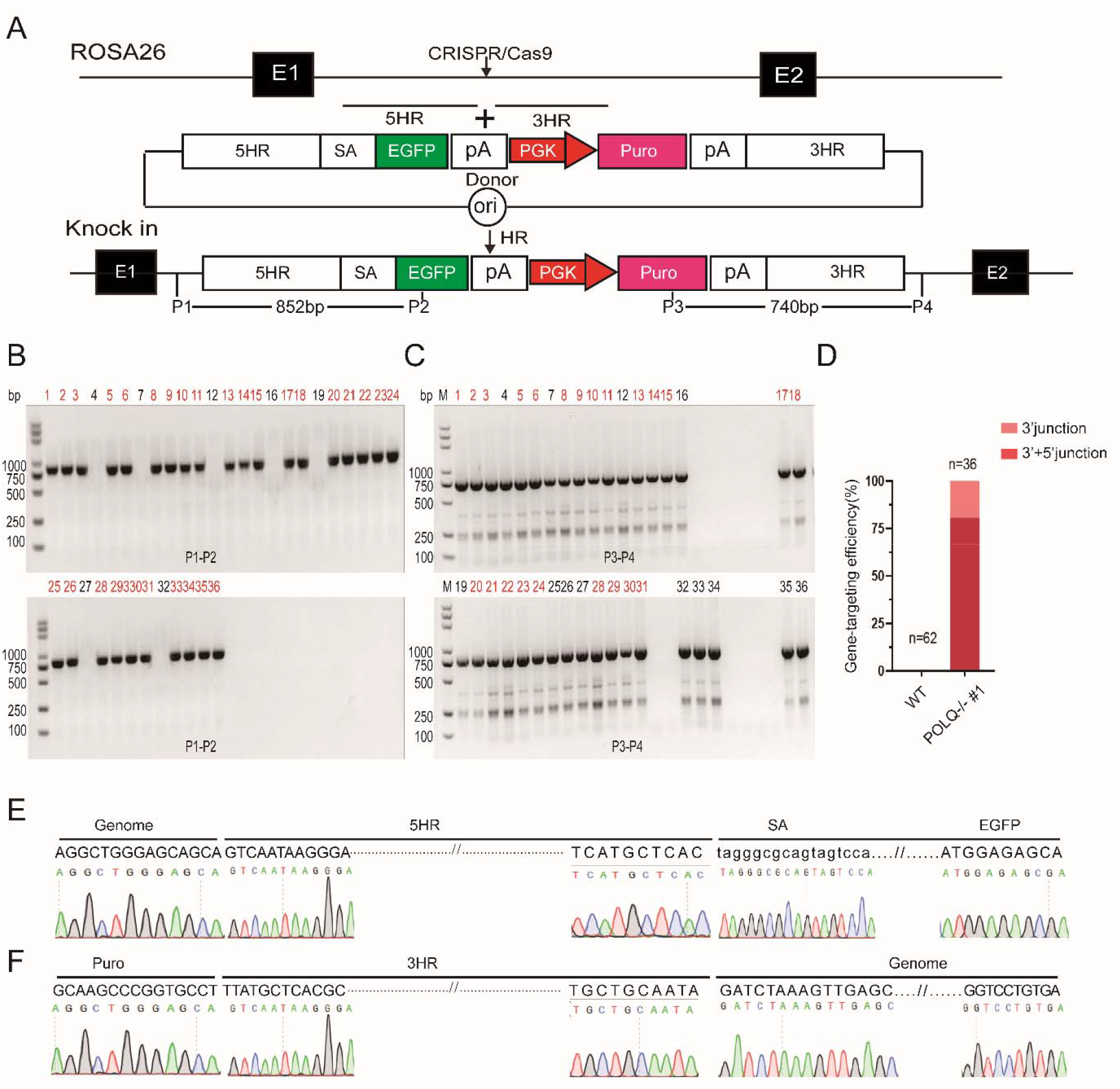
HR frequency in *POLQ*^−/−^ CHO cells. **(A)** A diagram for CRISPR/Cas9-mediated HR of Rosa26-EGFP-puro into the *Rosa26* locus in *POLQ*^−/−^ CHO cells. eGFP, enhanced green fluorescent protein gene; PGK, PGK promoter; Puro, Puromycin-resistance gene; SA, splice acceptor. **(B)** 5′ junction PCR analysis of the gene knock-in cell clones. M, 1 kb DNA ladder; Lanes 2-15, cell clones; WT, wild-type CHO cells; P, the donor vector; H2O was the negative control. **(C)** 3′ junction PCR analysis of the gene knock-in cell clones. **(D)** Gene-targeting efficiency of GFP in WT, *POLQ*^−/−^ CHO-KI cells, data shown are the mean ± SEM (n = 3). **(E)** Sequencing confirmation of the 5′ junction after targeted integration of Rosa26-EGFP-puro cassettes into the CHO *Rosa26* locus. **(F)** Sequencing confirmation of the 3′ junction after targeted integration of Rosa26-EGFP-puro cassettes into the CHO *Rosa26* locus.

### Loss of Pol θ inhibit random integration in CHO cells

To further investigate the reason why knockout *POLQ* improves HR efficiency in CHO cells, we investigated the involvement of *POLQ* in RI. We measured the ability of *POLQ* knockout CHO cells to form stable puromycin-resistant colonies upon RI of transfected plasmid DNA encoding a puromycin resistance gene (**Figure 4A**). Similar to previous results, we found fewer cell clones in *POLQ*^-/-^ cells (**Figure 4B**) and nearly 10-fold decrease in random integration efficiency in *POLQ*^-/-^ cells compared to WT (**Figure 4C**). This may suggest that Pol θ is important for the RI in CHO cells which is consist with previously research in mES and Nalm-6 cells.

**Figure 4.**
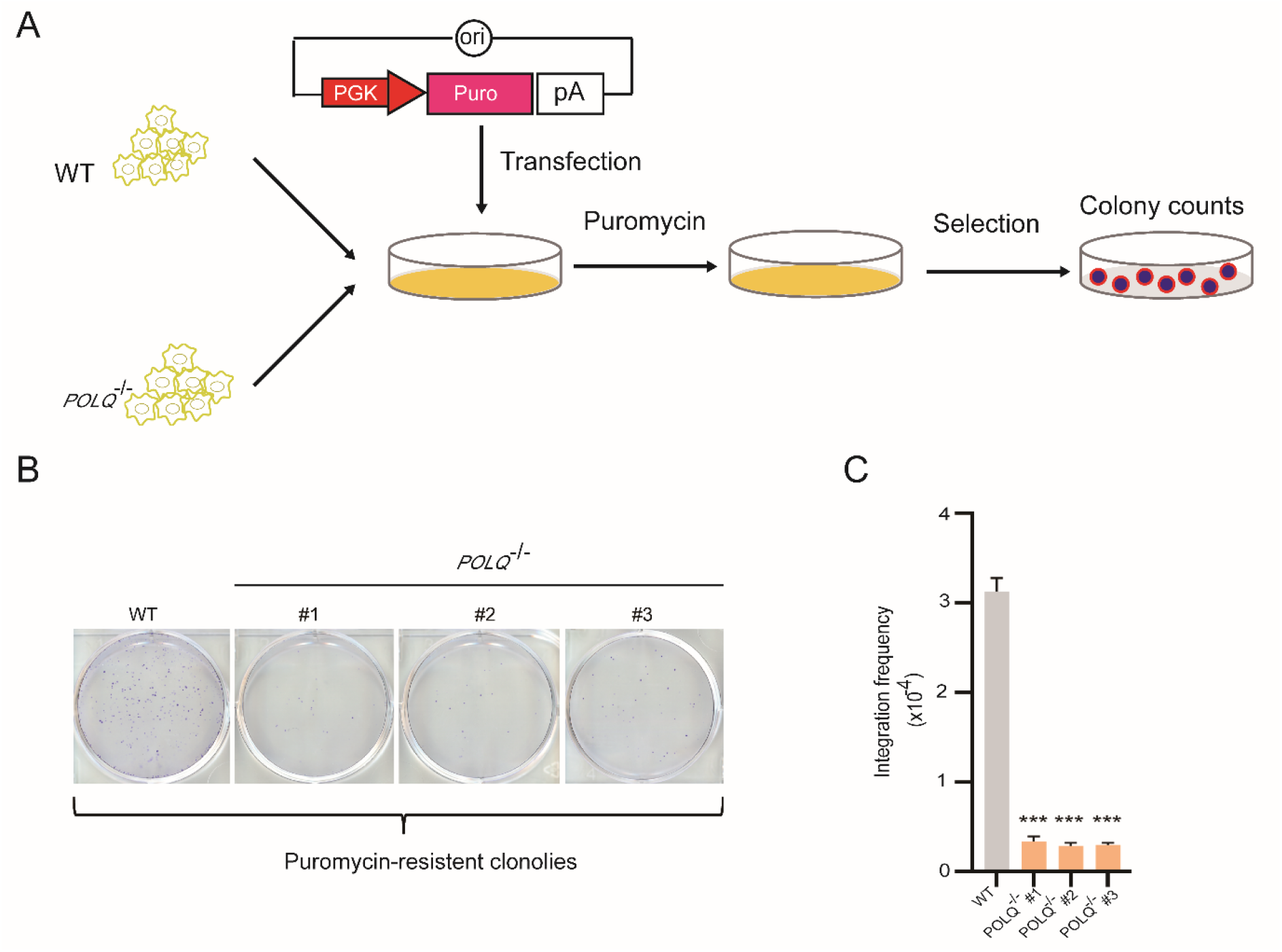
RI was suppressed in *POLQ*^−/−^ CHO cells. **(A)** Schematic of RI assay. PGK, PGK promoter; Puro, Puromycin-resistance gene. PGK-puro was electroporated into WT, *POLQ*^−/−^ cell lines, after three weeks with puromycin, colony formation were counted. **(B)** Representative colonies arose in WT, *POLQ*^−/−^ cell lines. **(C)** RI frequency in the WT, *POLQ*^−/−^ cell lines, data shown are the mean ± SEM (n = 3).

### RNA-sequence analysis reveals not significant altered DNA repair, metabolism, apoptosis, and cell cycle in *POLQ*^-/-^ cells

To get a global overview of the molecular pathways regulated by *POLQ* in CHO cells, we performed RNA-Sequence to identify gene expression alterations. According to only two groups (*POLQ*^-/-^ and WT) were being analyzed, we mapped a Venn diagram of gene expression, displaying the number of shared and exclusively expressed genes between the two groups. There were 18445 and 18256 expressed genes detected from the *POLQ*^-/-^ and WT group, respectively. Among them, 17665 were shared, 780 were exclusive to the *POLQ*^-/-^ group, and 591 were present only in the WT group (**Figure 5A**). Heat map was created to visualize the quantitative differences in the expression levels of the DEGs between the *POLQ*^-/-^ and WT group (**Figure 5B**). The heat map was performed based on Z scaled RNA-seq FPKM (expected number of Fragments Per Kilobase of transcript sequence per Millions base pairs sequenced) of RNA-seq data and represents the gene expression levels. The red region indicates high expression levels. The blue region indicates low expression levels. Colors change from red to blue, showing that the values of Z scaled FPKM change from high to low. In total, 335 significant differentially expressed genes (DEGs) were identified between the *POLQ*^-/-^ and WT group with DEG-seq analysis. 176 genes were up-regulated and 160 were down-regulated (**Figure 5C**). We performed pathway enrichment analysis on the genes of each module separately, explored their related functions, and obtained many meaningful KEGG pathway and GO terms (**Figure 5D, 5E**). In the KEGG enrichment analysis, it is significantly enriched to ECM-receptor interaction and PI3K-Akt signaling pathway. While we also obtained the pathways of metabolism, apoptosis, DNA repair, and cell cycle, but all these pathways were not statistically significant. All DEGs were classified into three GO categories by the “enrichGO” in “clusterProfiler”, including biological process (BP), cellular component (CC), and molecular function (MF). Among GO terms, only enzyme regulator activity and molecular function regulator of molecular function (MF) were significantly enriched. Meanwhile, we also counted the GO terms associated with DNA repair, metabolism, apoptosis, and cell cycle, and found that all of them were not statistically significant.

**Figure 5.**
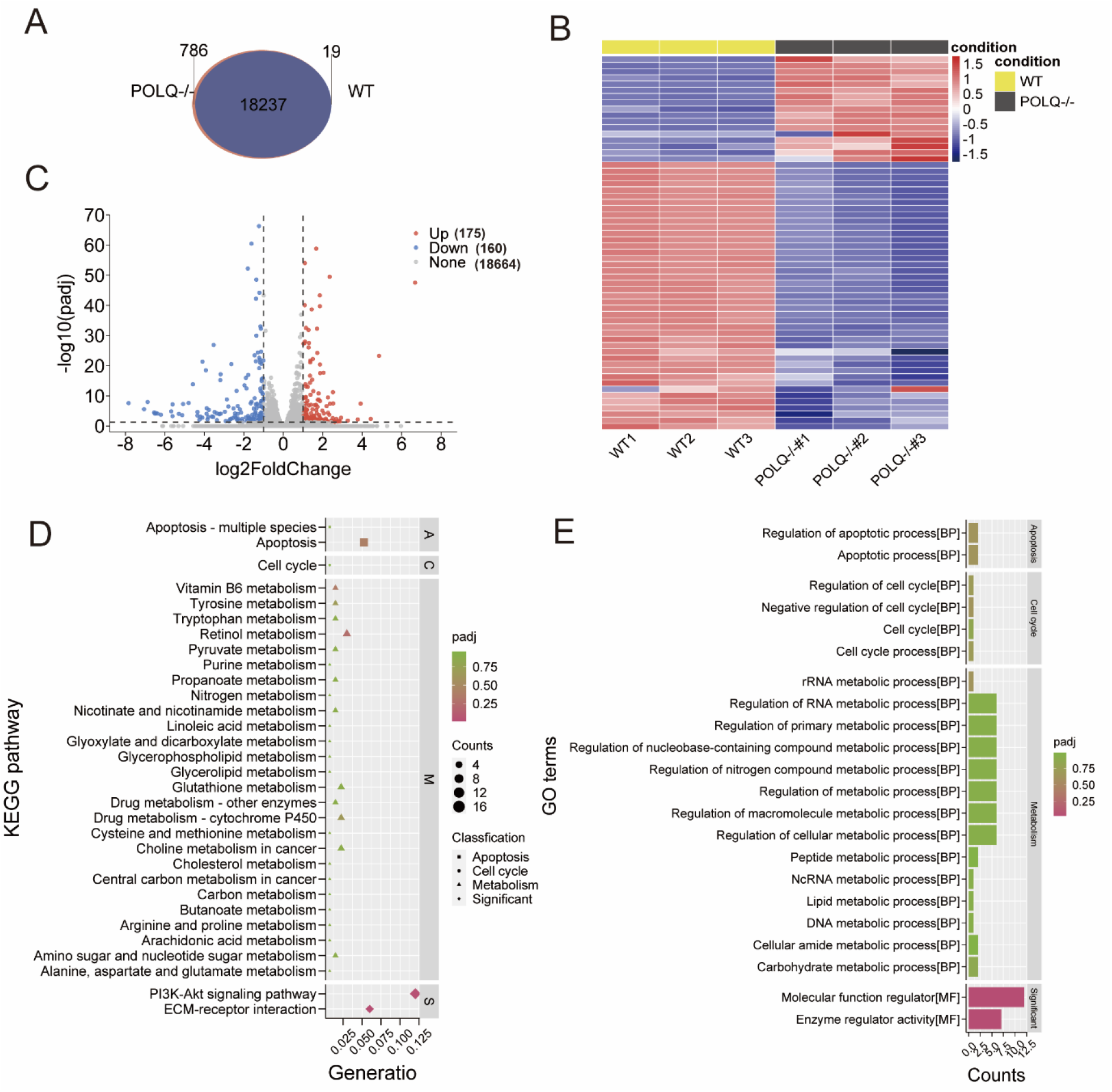
The Transcriptome analysis of the genes differentially expressed in *POLQ*^−/−^ and WT CHO cells. **(A)** Venn diagram shows differentially expressed genes in the *POLQ*^−/−^ and WT cells. **(B)** Heatmap shows the top 60 highly variant genes (x-axis: samples of WT and *POLQ*^−/−^; in red: upregulation, in blue: downregulation). **(C)** Volcano plot shows the differential expression analysis of the comparison *POLQ*^−/−^ vs. WT (genes above thresholds highlighted by coloring). **(D)** Bubble charts for KEGG pathway enrichment analysis of DEGs. Bubble size represents the gene ratio between the genes connected to a process and the overall query size. The color scale shows the padj value; **(E)** Bar charts for Gene Ontology.

## DISCUSSION

Here we show that manipulation of DNA repair pathway choice is an efficient strategy to maximize precision genome editing in CHO cells with CRISPR-Cas9. We reported that loss of Pol θ, which is important gene for Alt-EJ DNA repair pathway, can increases the efficiency of Cas9-stimulated HDR up to 30-fold in CHO cells by eGFP reporter system. Meanwhile, using a simple circular plasmid without negative selection element as the homologous template DNA donor, the two genomic sites in the *POLQ* knockout cell line can still efficiently obtain gene knock-in positive clones. The efficiencies are respectively 80% (25/36) and 2.7% (1/36), however no positive clones were obtained in wild-type cells. Furthermore, we show that Pol θ promotes random integration and RNA-sequence analysis reveals not significant altered DNA repair, metabolism, apoptosis, and cell cycle in CHO cells. Our research not only plays an important role in the precise gene editing of CHO cells, but also provides important information for the study of CHO cell DNA repair mechanisms.

Many previous studies inhibit C-NHEJ pathway by gene knockout (30), RNAi (31), proteolytic degradation (32) and small molecules (32-34) to improve the efficiency of HDR. However, the C-NHEJ pathway is important for the genome stability and inhibition of NHEJ can induce apoptosis (71). Meanwhile, if to inhibit of another Alt-EJ pathway can improve HR has poorly been reported. Recently, two research have been reported that dual loss of *POLQ* and *LIG4* abolishes random integration in mES (59) and Nalm-6 cells (58). However, inactivated the *POLQ* did not improve the efficiency of HDR in mES (59). Here, we found that knock out the *POLQ* can robustly improve the efficiency of HDR in CHO cells. This may be the cell type dependent manner such as the effect of scr7 in different cells (46,47,52-54). Furthermore, Pol θ is a unique DNA polymerase in that it is highly expressed in various types of cancer (72,73). Intriguingly, loss of Pol θ is synthetically lethal with HR defects and thus developing Pol θ inhibitors will lead to a new therapeutic agent for various cancers with compromised HR (56,57). It has been reported that *POLQ* only plays a minor role in repair of DSBs compare with *LIG4* (58). So, inhibition of *POLQ* is a better target gene selection.

Furthermore, it has been reported that coupled the use of CRISPR/Cas9 with homology-directed repair (HDR) for targeted integration of homologous donors in CHO cells (54,74). However, the frequency of HR-mediated genome editing in CHO cells is very low with the rate ranging from 1% to 27.8%. This makes isolation of clones with targeted gene insertions time-consuming and labor intensive, creating a desire for more efficient approaches to accelerate genome engineering. It has been reported that fluorescent enrichment resulted strategy in a threefold increase in the number of cells with HDR-mediated genome editing at the two target sites relative to non-enriched samples, yielding HDR frequencies of 7% by transient setup. But, only 1% HDR frequencies was obtained by screening of knock-in CHO cells clones (54). Additionally, optimize the donor vector can also improve the HDR frequencies to 13%-18% (75), 26%-36% (51) respectively by transient setup and 25.4∼66.0% (76) by screening of knock-in CHO cells clones. Recently, knockdown Mre11 and Pari, and elevated Rad51 expression levels can improve precise genome editing up to 75% in CHO cells by transient setup (77). To improve and simple the stage, we only knockout the *POLQ* gene. Two strategies were employed to measure the CRISPR/Cas9-mediated HDR efficiency in our research. One strategy is an eGFP reporter system to detect. Results from the eGFP reporter system demonstrated loss of Pol θ led to a 30-fold increase in HDR efficiency. Another strategy is the screening of knock-in CHO clones in two genome sites as the *Rosa26* and *COSMC* to detect the HDR efficiency. Meanwhile, we used a simpler circular plasmid without negative selection element as the donor vector, which simplified the size and operation of the donor vector (51,54,74). The knock-in rate of 4 kb simple circular DNA donor integration can reach a mean of over 80% (25/36) and 2.7% (1/36) of the selected cell colonies in *POLQ*^-/-^ CHO cells which is higher than previously report (74). However, no positive knock-in cell colonies were obtained in WT CHO cells which respectively selected 62 cell colonies and 36 cell colonies.

Recently, it has been reported that efficient HR-mediated genome editing depends on Alt-EJ DNA repair activities (77) which is contradicts our research in CHO cells and previous research on other cells (58,59). This may be due to the different technology used, they use RNAi technology (77). Although our technology has been successful in CHO cells, it is not suitable for all types of cells, such as large animal primary cells, because gene knockout is lethal. After editing large animal cells, nuclear transfer is often performed to obtain surviving animals. The latest CAS9i technology or small molecular inhibitor may be a solution (31).

Although, it has been reported that *POLQ* only plays a minor role in repair of DSBs. We also performed RNA-Sequence to get a global overview of the molecular pathways regulated by *POLQ* in CHO cells and found that the GO terms associated with DNA repair, metabolism, apoptosis, and cell cycle, and found that all of them were not statistically significant. Specially, DNA repair path way related genes such as Ku70, Ku80, LIGI, LIGIII and RAD51 were not significant altered which is consist with the previously result that POLQ may prevent the assembly of RAD51 monomers into RAD51 polymers not alter the expression of the RAD51.

Although we did not detect off-target mutations in any of the potential off-target sites in our experiments using DNA sequencing and T7E1 assay, however, off-target effects could not be ruled out. Therefore, the whole-genome sequencing technology can be used for off-target detection in later studies. Meanwhile, ribonucleoproteins (RNPs), the Cas9 protein in complex with in vitro transcribed guide RNA, have been used together with a single-stranded oligonucleotide HDR template for nucleotide replacement or insertion (45,78,79). We believe that our technology can also be delivered together with Cas9 protein, sgRNA transcripts and the double cut donor (49,75,76), which may even be preferable in industrial applications.

Taken together, the present study describes a new strategy for the enhancement of HDR in CHO cells. DSB repair pathway choice can be redirected by inhibiting Alt-EJ pathway via knockout *POL*Q, shifting the HDR/NHEJ balance several folds. These findings provide an efficient approach to promote precise gene editing in mammals’ cells, supporting its applications for research and therapy.

## Materials and methods

### Vector construction

To construct the CRISPR/Cas9 expression vectors, candidate sgRNAs were designed using online design tool (http://crispr.mit.edu/), and each 20-bp target sequence was subcloned into the pX330 vector (Addgene 42230). The CRISPR/Cas9 target sequences (20-bp target and 3-bp PAM sequence) used in this study are listed in **Table S1**. The pPGKneoDTA vector was used as the backbone to construct the *Rosa26* gene-targeting vector. A 843 bp 5’ homologous arm and a 910 bp 3’ homologous arm were amplified by PCR from the genome of CHO-KI, and cloned into the pPGKneoDTA vector. A splice acceptor (SA) and a promoterless eGFP (SA-EGFP) element and a puromycin expression cassette (PGK-promoter, puromycin, BGH pA) were synthesized by Sangon Biotech. Conventional molecular cloning and Golden Gate assembly were used to construct pROSA26-EGFP, pROSA26-EGFP-puro vector.

### Cell culture and transfection

The CHO-K1 cells (Invitrogen, Carlsbad, CA, USA) were cultured in Dulbecco’s modified Eagle’s medium (DMEM; Gibco, Grand Island, New York, USA) supplemented with 10% fetal bovine serum (FBS; Gibco, Grand Island, New York, USA) at 37.5°C in an atmosphere of 5% CO2 and humidified air. CHO-K1 cells were thawed and cultured for two days until reaching subconfluence before transfection. DNA transfection using an Lipofectamine 3000 reagent (Invitrogen, Carlsbad, CA, USA). Briefly, cells were washed twice with Dulbecco’s Phosphate Buffered Saline (DPBS; Gibco, Grand Island, New York, USA) and an aliquot of the cell suspension (1 × 10^6^ cells) was transfected with 3 μg different donor plasmids and/or sgRNA plasmids. Following transfection, cells were then cultured for 24 h and replated for cell colonies forming.

### T7EI assay

The editing activity of each Cas9 vector was assayed using T7 endonucleaseI (T7EI) (New England Biolabs, Ipswich, MA, USA) as described previously. Briefly, genomic DNA from Cas9-treated cells was extracted using a DNeasy Blood and Tissue kit (QIAGEN, Hilden, Germany). PCR amplicons including nuclease target sites were generated using the primer pairs POLQ-F/POLQ-F for the *POLQ* locus. The primers are listed in **Table S2**. Six hundred nanograms of purified PCR product was annealed according to the following settings: 95 °C for 5 min and ramp down to 25 °C (at 0.5 °C/min). The annealed product was then digested with T7EI at 37 °C for 30 min and separated on 2% agarose gel. Mutation frequencies (% indels) were calculated by measuring band intensities with ImageJ software. The band intensities were quantified based on relative band intensities.

### Generation of gene-knockout cell lines

For the generation of *POLQ*-knockout cell lines, limiting dilution was used. Briefly, the single cell was separated by fluorescence-activated cell sorting (FACS) 24-48 h after transfection, and transferred to 96-well plates with 10% FBS containing puromycin (10 μg/mL) for 3 weeks. After selection, expanded, and cryopreserved, the colonies were analyzed. PCR amplicons including nuclease target sites were generated using the primer pairs POLQ-F/POLQ-F (**Table S2**). PCR-products was used to identify clones with a bi-allelic mutation. PCR-products of clones that showed bi-allelic mutation were sent for sequencing to confirm the introduction of a premature stop-codon due to deletions/insertions that cause a frameshift.

### Western blotting

Samples were isolated from the gene-knockout cell lines and wild-type (WT) CHO-KI cells, and homogenized in cell lysis buffer for Western and IP analyses (Beyotime, Shanghai, China). After centrifugation at 10000 g for 10 min at 4 °C, the total protein supernatants were collected, and protein concentrations were measured using a BCA Protein Assay kit (Beyotime, Shanghai, China). Approximately 50 μg of protein was separated on 7.5% SDS-PAGE gels and transferred to Immobilon-P membranes (MilliporeSigma, Burlington, MA, USA). After blocking in 5% milk, 0.05% Tween 20 PBS for 1 h, membranes were incubated with a POLQ antibody (1:10000; Abcam, Cambridge, MA, USA), β-actin antibody (1:10000; Abcam, Cambridge, MA, USA) was used as a reference. After washes with PBST, membranes were incubated with a goat anti-rabbit antibody conjugated with horseradish peroxidase (1:20000; Sino-American Co, Beijing, China) for 1 h followed by three washes with PBST. Protein signals were detected using an ECL Chemiluminescence kit (Thermo Fisher Scientific, Waltham, MA, USA).

### Determination of Homologous Recombination Efficiency

To determine HR efficiency, wild-type and knockout cells were seeded into six-well plates and transfected with ROSA26-AdSA-TurboGFP-polyA plasmids and sgRNA-Cas9 vectors targeting *ROSA26*. After 72 h, GFP+ cells were photographed under a fluorescence microscope (Olympus IX83, Tokyo, Japan). Then, digested cells were resuspended in FACS buffer (2% FBS–PBS) and analyzed by a FACS Calibur flow cytometer (Becton, Dickinson and Company, USA). Compensation and analysis of results were performed using Flow Jo software.

### Random integration assay and gene-targeting assays

For Random integration (RI) assays, pPGK-puro was linearized and transfected into *POLQ*^−/−^, and wild-type (WT) CHO-KI cells. After cultivation for 2–3 weeks, the resulting puro-resistant colonies were counted and the total integration frequency was calculated by dividing the number of puro-resistant colonies with that of surviving cells. For gene-targeting assays, linearized pROSA26-EGFP and sgRNA were transfected into *POLQ*^−/−^ and WT cells. After a 72 h incubation, cells were trypsinized, collected by centrifugation, and resuspended in 1 ml PBS, GFP^+^ cells were analysed by FACS using BD LSR Fortessa instrument (BD, New York, USA).

### Junction analysis of gene-targeting clones

The pROSA26-EGFP-puro targeting vector and sgRNA were cotransfected into *POLQ*^−/−^ and WT CHO-KI cells. Cells were seeded at low density 48 h after transfection and maintained with 10% FBS containing puromycin (10 μg/mL) for 8–10 days until colonies were formed. Colonies were picked and grown in 96-wells format, at (semi) confluence cells were split to two 96-wells plates. One plate was used for sub-culturing of cells; the other plate was used for DNA isolation and 5′/3′ junction PCR analysis of the individual clones. The P1/P2 primer pair was used for the 5’ arm and produced a 2.7-kb amplicon, and the P3/P4 primer pair was used for the 3’ arm and produced a 2.2-kb amplicon (**Table S2**), to confirm the successfully gene-targeting cell clones. PCR was performed for 35 cycles at 94 °C for 30 s, 60 °C for 30 s, and 72 °C for 2-3 min with a hold at 72 °C for 10 min. Purified 5′/3′ junction PCR product was sequenced by Sangon Biotech.

### RNA extractions from cultivations

Samples for RNA extractions were taken from *POLQ*^−/−^ and WT CHO-KI cells. The withdrawn sample was immediately cooled on ice and the pellet was harvested by centrifugation at 4°C, washed with cold water and the biomass stored at −80°C until further treatment. Total RNAs were extracted from *POLQ*^−/−^ and WT CHO-KI cells using Trizol (Invitrogen) followed by additional purification using an RNeasy Mini Kit (QIAGEN, Hilden, Germany) according to the manufacturer’s instruction. The quality of the RNA was assayed using a BioAnalyzer (Agilent Technologies, Palo Alto, CA, USA). The high quality RNAs were used for constructing the library that was used for sequencing.

### Pre-processing and quality assurance of the RNA-seq reads

For RNA-seq analysis, Illumina NovaSeq 6000 was used to perform paired-end sequencing of the RNA samples using the standard Illumina RNA-seq protocol with a pair-end 150 bp reads. All raw RNA-seq data generated in this study will be available. The raw reads form RNA were processed quality control (QC) with in-house perl scripts. Bad quality reads containing adapter sequence, contaminant DNA and low-quality reads were removed, and clean data were generated. Clean reads were then indexed and mapped to reference genome (60) of CHO-KI cell using Hisat2 (v2.0.5) (61). The aligned records from the aligners in BAM/SAM format (62) were further examined for potential duplicate molecules in each record and removed using the Picard tool kit (63). After that, gene expression levels were estimated using FPKM values by the featureCounts (v1.5.0-p3) (64) software.

### Identification of differential gene expression

The differential gene expression (DEGs) between *POLQ*^−/−^ and WT CHO-KI condition were identified using the DESeq2 R package (65). The resulting P-values were adjusted using the Benjamini and Hochberg’s approach for controlling the false discovery rate. padj<0.05 and |log2(foldchange)| > 1 were set as the threshold for significantly differential expression.

### Gene ontology enrichment analysis

Gene Ontology (GO) and Kyoto Encyclopedia of Genes and Genomes (KEGG) pathway enrichment analyses were performed to predict the potential function of differentially expressed genes (DEGs) by R (4.1.0)”ClusterProfiler” package (66). GO terms and KEGG pathway with corrected P-value < 0.05 and false discovery rate (FDR) < 0.05 were considered statistically significant. All statistical analyses and illustrations were done under R suite software.

### Statistical Analyses

Data were analyzed using unpaired two-tailed Student’s *t*-tests or one-way ANOVA followed by Bonferroni’s post-hoc test for multiple groups. Figures were prepared using GraphPad Prism v8.4 (GraphPad Software, San Diego, CA, USA). Error bars represent the Standard Error of Mean (SEM).

## DATA AVAILABILITY

All data in the manuscript or the supplementary material is available upon request. The code and sample dataset of ERMer are available in the Github repository (https://github.com/tibbdc/ermer).

## ACKNOWLEDGEMENTS

We thank Ke Zhao (Beijing Institute of Lifeomics, China) for imaging and Ming Yu (Beijing Institute of Pharmacology and Toxicology, China) for FACS.

## AUTHOR CONTRIBUTIONS

J.W., and Z.S. conceived the project and designed the experiments. C.W., M.W., Y.W., X.L., C.L., Y.Z. performed the experiments. M.Z. contributed to bioinformatic analysis. C.W., M.W., Y.W. did the data analysis. B.S., J.W.and J.F. supervised the project. J.W. and Z.S. wrote the manuscript. All authors discussed and approved the manuscript.

## SUPPLEMENTARY INFORMATION

## Supplementary Figure S1-3

**Figure S1.**
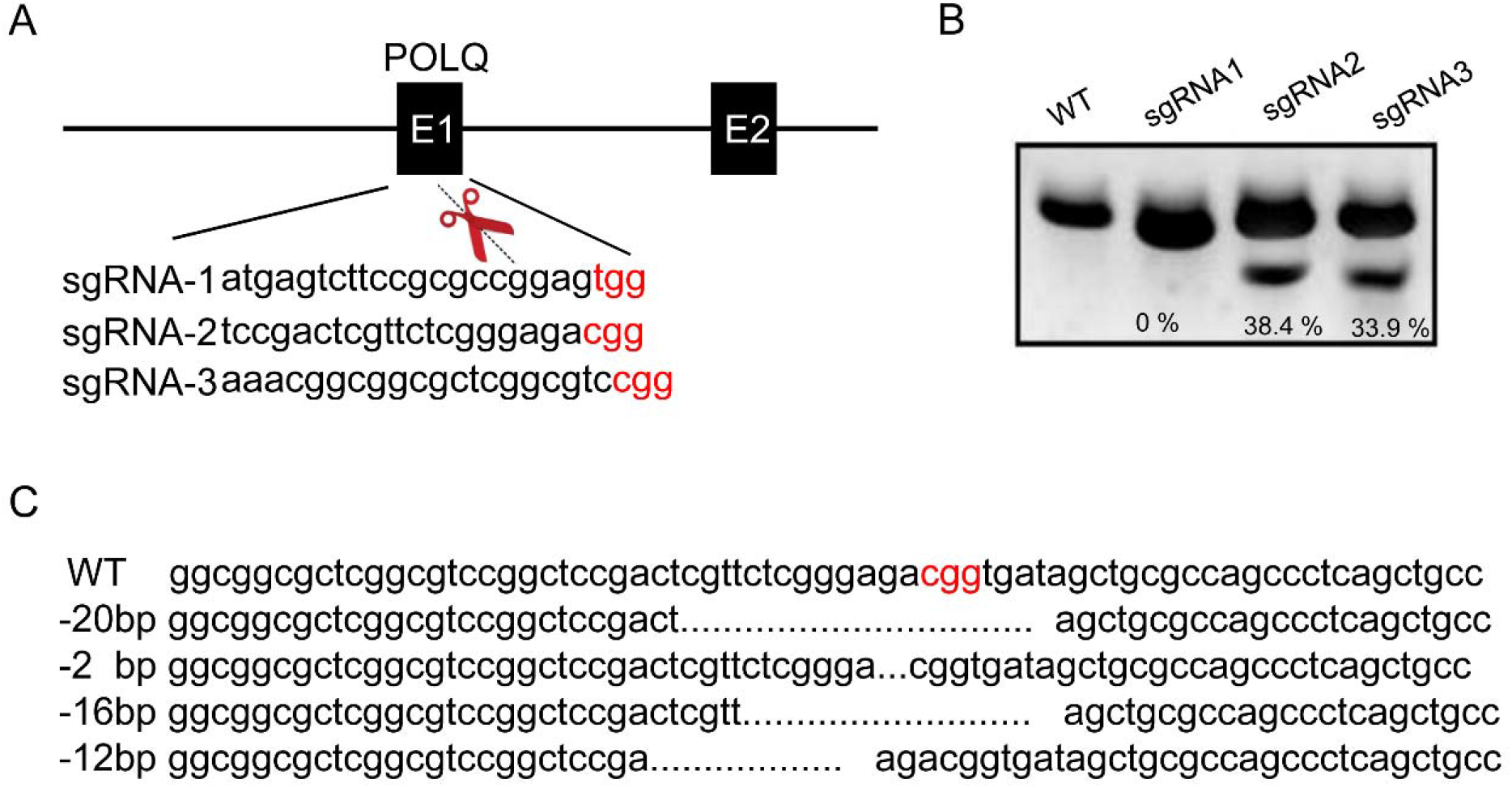
Design of Cas9 for the induction of DSBs in the endogenous *POLQ*. **(A)** Modification of the CHO *POLQ* gene by CRISPR/Cas9. The CRISPR/Cas9 target sequences (20-bp target and 3-bp PAM sequence (colored in red)) are shown. **(B)** Representative results of T7EI assays of sgRNAs directed against the CHO *POLQ*. The mutation frequencies (% indels) of different sgRNAs were calculated by measuring the band intensities. WT, wild-type cells; sgRNA-1-3, sgRNAs against the CHO *POLQ* transfected cells; **(C)** Representative sequencing results of TA clones revealing different indel mutations mediated by sgRNA-2 in the *POLQ* target site.

**Figure S2.**
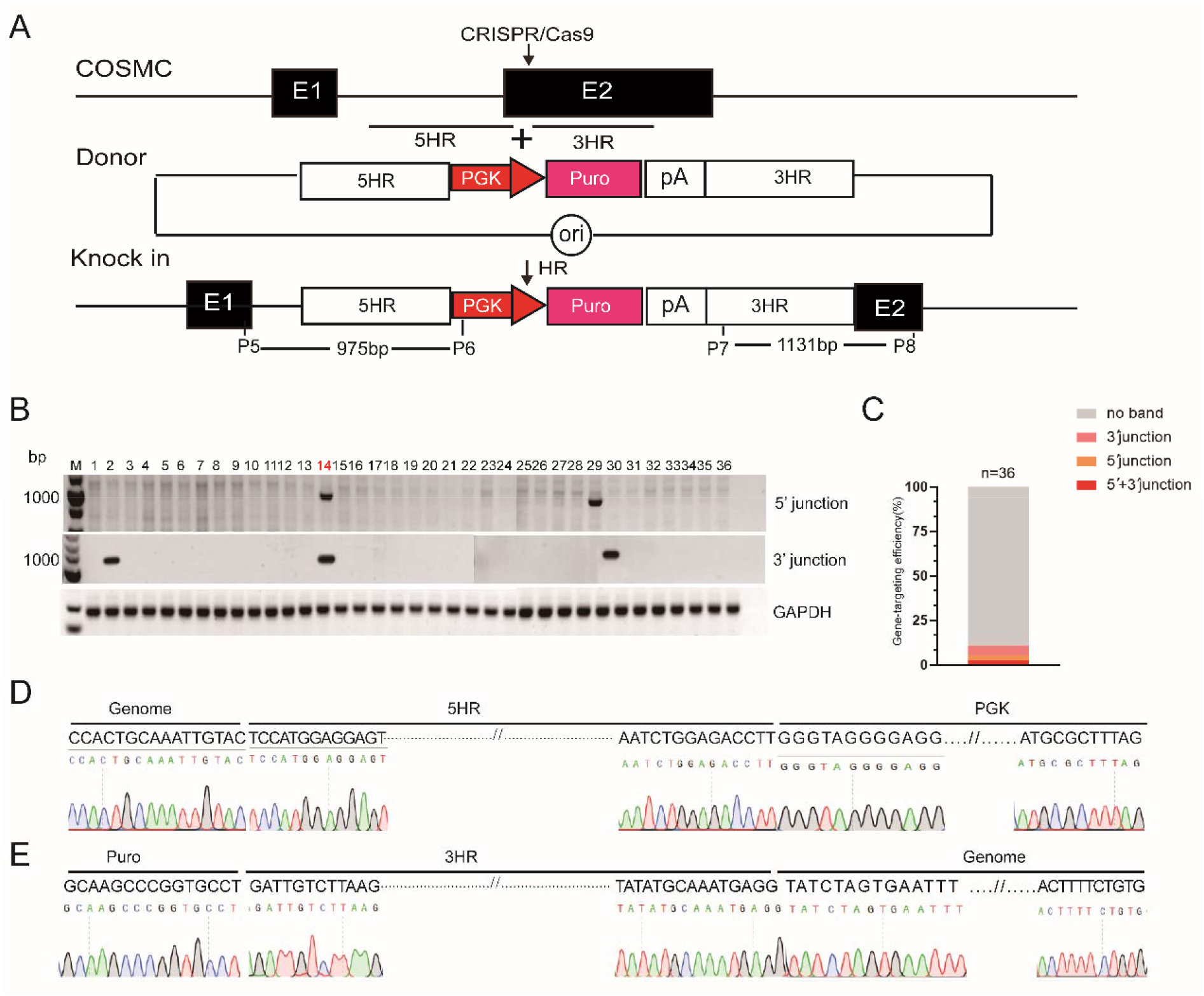
HR frequency in *POLQ*^−/−^ CHO cells. **(A)** A diagram for CRISPR/Cas9-mediated HR of PGK-Puro into the COSMC locus in *POLQ*^−/−^ CHO cells. PGK, PGK promoter; Puro, Puromycin-resistance gene. **(B)** 5′ junction and 3′ junction PCR analysis of the gene knock-in cell clones. M, 1 kb DNA ladder; Lanes 2-15, cell clones; WT, wild-type CHO cells; P, the donor vector; H2O was the negative control. **(C)** Gene-targeting efficiency of Puro in WT, *POLQ*^−/−^ CHO-KI cells, data shown are the mean ± SEM (n = 3). **(D)** Sequencing confirmation of the 5′ junction after targeted integration of PGK-puro cassettes into the CHO COMSC locus. **(E)** Sequencing confirmation of the 3′ junction after targeted integration of PGK-puro cassettes into the CHO COMSC locus.

## Supplementary Table S1-2

**Table S1.**
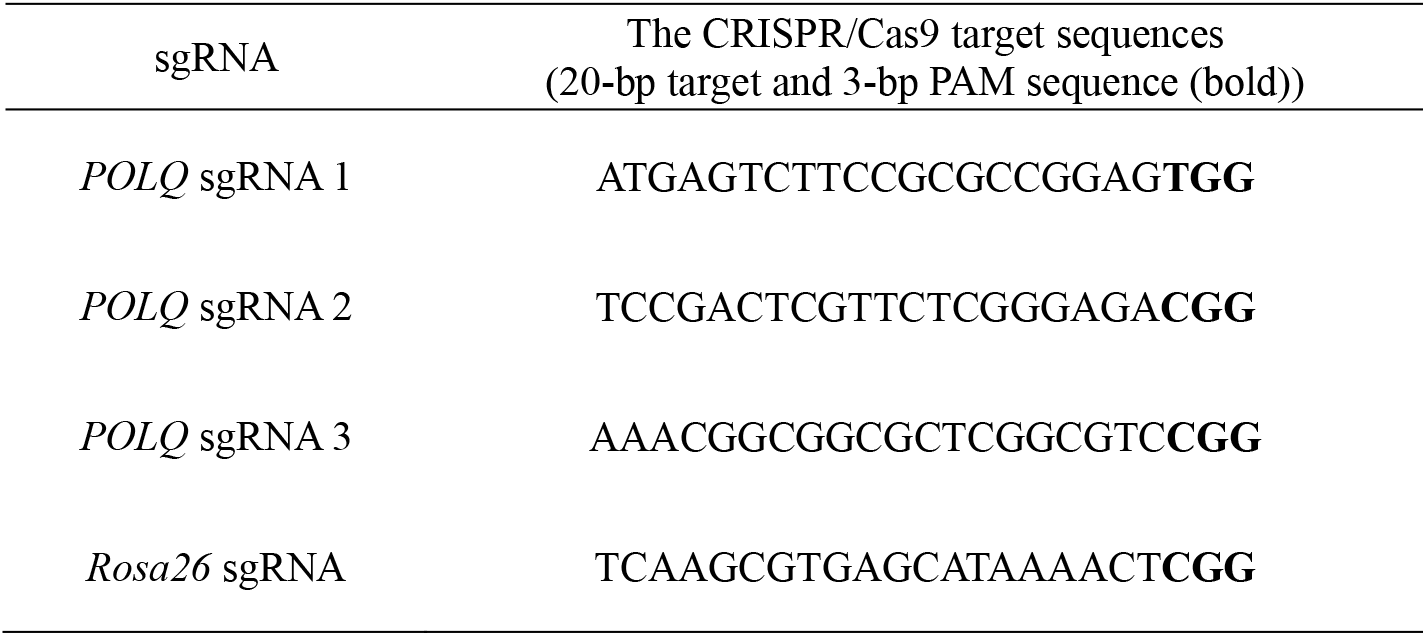
Target sequences of the sgRNAs used in this study.

**Table S2.**
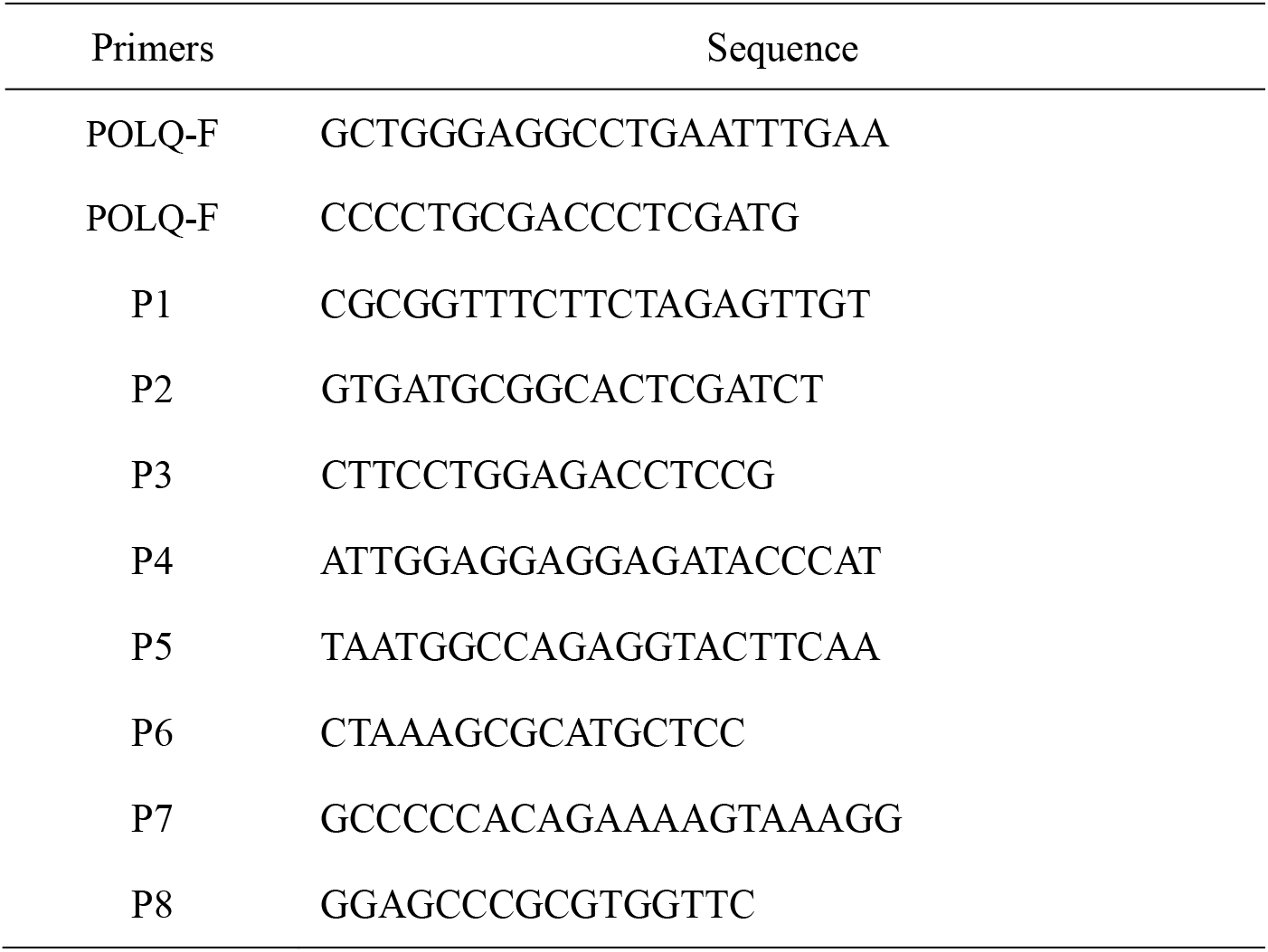
List of primers used in the present study.

