## Supplementary Files for "Inhibiting POLQ-mediated alternative NHEJ enhances CRISPR/Cas9 mediated precise genome editing in CHO cells"

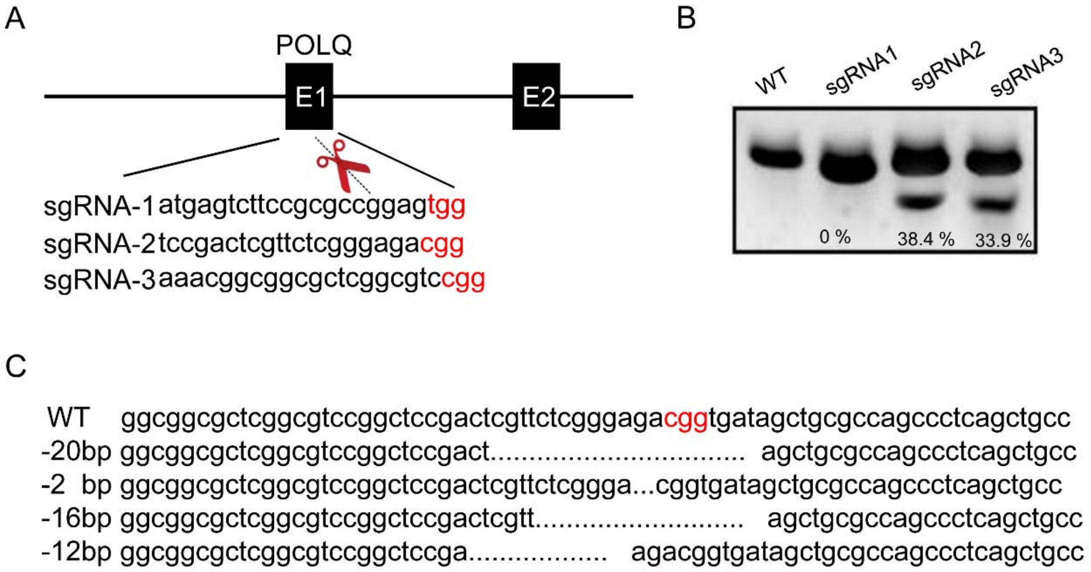


**Figure S1. Design of Cas9 for the induction of DSBs in the endogenous *POLQ*. (A)** Modification of the CHO *POLQ* gene by CRISPR/Cas9. The CRISPR/Cas9 target sequences (20-bp target and 3-bp PAM sequence (colored in red)) are shown. **(B)** Representative results of T7EI assays of sgRNAs directed against the CHO *POLQ*. The mutation frequencies (% indels) of different sgRNAs were calculated by measuring the band intensities. WT, wild-type cells; sgRNA-1-3, sgRNAs against the CHO *POLQ* transfected cells; **(C)** Representative sequencing results of TA clones revealing different indel mutations mediated by sgRNA-2 in the *POLQ* target site.

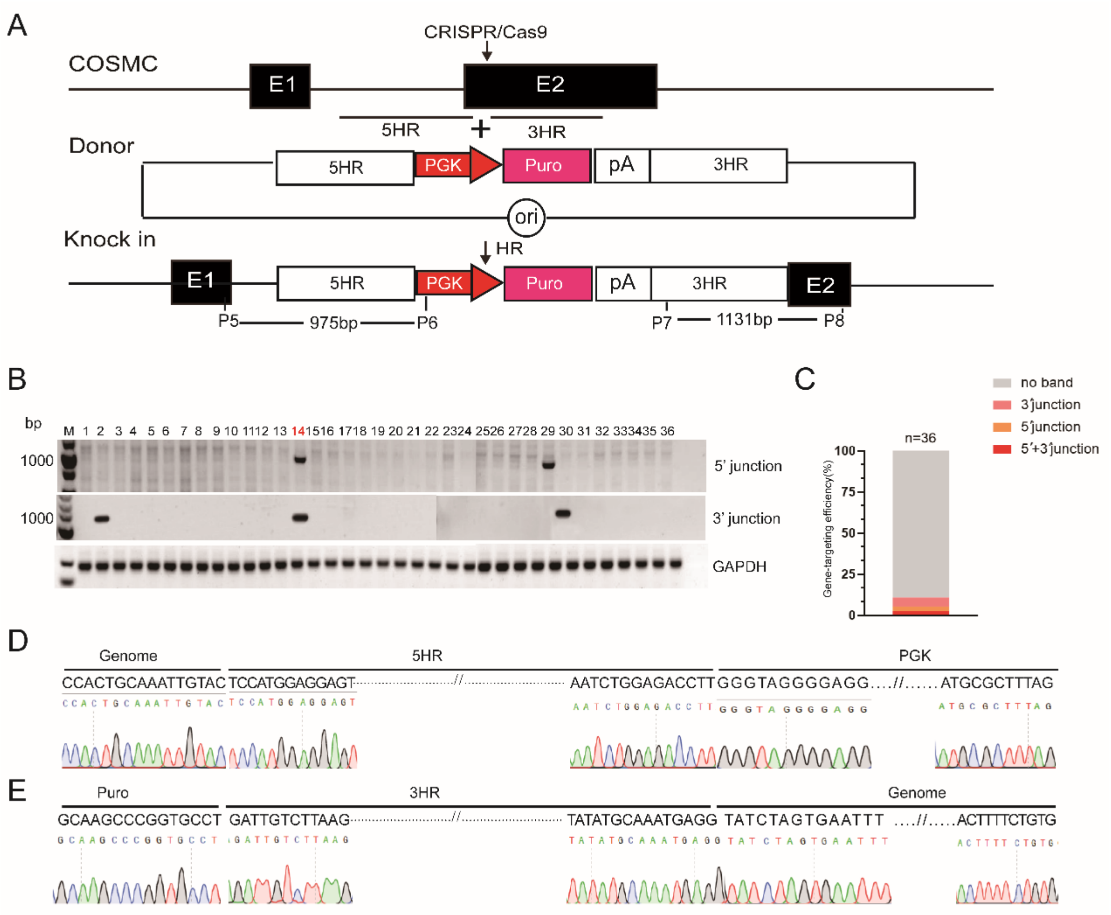


**Figure S2. HR frequency in *POLQ*$^{-/-}$ CHO cells. (A)** A diagram for CRISPR/Cas9-mediated HR of PGK-Puro into the COSMC locus in *POLQ*$^{-/-}$ CHO cells. PGK, PGK promoter; Puro, Puromycin-resistance gene. **(B)** 5′ junction and 3′ junction PCR analysis of the gene knock-in cell clones. M, 1 kb DNA ladder; Lanes 2-15, cell clones; WT, wild-type CHO cells; P, the donor vector; H2O was the negative control. **(C)** Gene-targeting efficiency of Puro in WT, *POLQ*$^{-/-}$ CHO-KI cells, data shown are the mean ± SEM (n = 3). **(D)** Sequencing confirmation of the 5′ junction after targeted integration of PGK-puro cassettes into the CHO COMSC locus. **(E)** Sequencing confirmation of the 3′ junction after targeted integration of PGK-puro cassettes into the CHO COMSC locus.

**Table S1 Target sequences of the sgRNAs used in this study**

| sgRNA | The CRISPR/Cas9 target sequences (20-bp target and 3-bp PAM sequence (bold)) |
|---|---|
| *POLQ* sgRNA 1 | ATGAGTCTTCCGCGCCGGAG**TGG** |
| *POLQ* sgRNA 2 | TCCGACTCGTTCTCGGGAGA**CGG** |
| *POLQ* sgRNA 3 | AAACGGCGGCGCTCGGCGTC**CGG** |
| *Rosa26* sgRNA | TCAAGCGTGAGCATAAAACT**CGG** |

**Table S2 List of primers used in the present study**

| Primers | Sequence |
|---|---|
| POLQ-F | GCTGGGAGGCCTGAATTTGAA |
| POLQ-F | CCCCTGCGACCCTCGATG |
| P1 | CGCGGTTTCTTCTAGAGTTGT |
| P2 | GTGATGCGGCACTCGATCT |
| P3 | CTTCCTGGAGACCTCCG |
| P4 | ATTGGAGGAGGAGATACCCAT |
| P5 | TAATGGCCAGAGGTACTTCAA |
| P6 | CTAAAGCGCATGCTCC |
| P7 | GCCCCCACAGAAAAGTAAAGG |
| P8 | GGAGCCCGCGTGGTTC |
